# Immunoglobulin fragment F(ab’)_2_ against RBD potently neutralizes SARS-CoV-2 in vitro

**DOI:** 10.1101/2020.04.07.029884

**Authors:** Xiaoyan Pan, Pengfei Zhou, Tiejiong Fan, Yan Wu, Jing Zhang, Xiaoyue Shi, Weijuan Shang, Lijuan Fang, Xiaming Jiang, Jian Shi, Yuan Sun, Shaojuan Zhao, Rui Gong, Ze Chen, Gengfu Xiao

**Affiliations:** State Key Laboratory of Virology, Wuhan Institute of Virology, Center for Biosafety Mega-Science, Chinese Academy of Sciences, Wuhan 430071, China; Wuhan YZY Biopharma Co., Ltd, Wuhan 430075, China; Shanghai Serum Bio-technology Co., Ltd, Shanghai 201701, China; University of the Chinese Academy of Sciences, Beijing 100039, China; CAS Key Laboratory of Special Pathogens and Biosafety, Wuhan Institute of Virology, Center for Biosafety Mega-Science, Chinese Academy of Sciences, Wuhan 430071, China

**Keywords:** SARS-CoV-2, neutralizing antibody, receptor binding domain, immunoglobulin fragment, COVID-19

## Abstract

COVID-19 caused by the emerging human coronavirus, SARS-CoV-2, has become a global pandemic, leading a serious threat to human health. So far, there is none vaccines or specific antiviral drugs approved for that. Therapeutic antibodies for SARS-CoV-2, was obtained from hyper immune equine plasma in this study. Herein, SARS-CoV-2 RBD with gram level were obtained through Chinese hamster ovary cells high-density fermentation. The binding of RBD to SARS-CoV-2 receptor, human ACE2, was verified and the efficacy of RBD in vivo was tested on mice and then on horses. As a result, RBD triggered high-titer neutralizing antibodies in vivo, and immunoglobulin fragment F(ab’)_2_ was prepared from horse antisera through removing Fc. Neutralization test demonstrated that RBD-specific F(ab’)_2_ inhibited SARS-CoV-2 with EC_50_ at 0.07 μg/ml, showing a potent inhibitory effect on SARS-CoV-2. These results highlights as RBD-specific F(ab’)_2_ as therapeutic candidate for SARS-CoV-2.

## 1 Introduction

In recent years, emerging or remerging viruses such as Severe Acute Respiratory Syndrome-coronavirus (SARS-CoV), Ebola, Lassa, Zika, H1N1 influenza, Middle East Respiratory Syndrome-coronavirus (MERS-CoV) and others, challenged the global biosafety system and attracted high attention from the world. SARS-CoV-2, firstly identified at Wuhan, China in 2020^1^, leading coronavirus disease 2019 (COVID-19), have caused a global pandemic of more than 1 million confirmed cases and 70 thousands deaths (average fatality >5%). Until now, most countries in the world are in peak outbreak and humans are suffering from the risk of SARS-CoV-2 infection. Unfortunately, no vaccines or drugs have been approved for clinical use, in spite of some already being in clinical trials^2, 3^, such as Chloroquine and Remdesivir^4, 5^. The epidemic situation urgently call for effective, specific and quickly accessible drugs^6^.

Neutralizing antibodies (nAbs) play important roles in antivirals^7–9^, benefiting from that they effectively inhibit viruses at entry stage, such as preventing viral attachment or membrane fusion. Polyclonal antibodies such as convalescent plasma from recovered patients were usually made as emergency treatments for emerging infectious diseases^10–13^. However, lack of blood source and risk of blood-borne diseases impede the wide clinical application of convalescent plasma^14^. Antisera produced by large animals like horses through passive immunization provides an alternative for that^15–17^. And the commercial process of obtaining horse antiserum and its derivatives is mature in the modern pharmaceutical industry.

As we know, SARS-CoV-2 was reported to employ angiotensin converting enzyme 2 (ACE2) to enter host cells^18^, using the same receptor with SARS-CoV^19^. Note that the amino acid sequence identity between SARS-CoV-2 and SARS-CoV spike proteins (S) is about 76% ^20^. The S protein was consisted of S1 and S2, of which, S1 is responsible for receptor attachment and S2 is responsible for membrane fusion^21^. The receptor binding domain (RBD) of SARS-CoV S1 could potently induce nAbs in vivo^22^, thus SARS-CoV-2 RBD theoretically can be a good immunogen to motivate nAbs in vivo.

Based on the above, SARS-CoV-2 RBD was expressed by mammalian cells, and its antigenicity and efficacy were tested both in mice and horses. Using traditional systemic immunization with an immunogen dose-increasing strategy, RBD elicited high-titer nAbs in horses. F(ab’)_2_ was acquired by removing Fc from IgG, and its efficacy was evaluated through a neutralization test on live virus in vitro. F(ab’)_2_ reported here validates the efficacy of RBD in triggering nAbs in vivo and is highlighted as an alternative to immunotherapy for COVID-19.

## 2 Results

### 2.1 Designation, preparation, and characterization of SARS-CoV-2 RBD

Aiming at effectively eliciting nAbs without triggering unrelated antibodies in vivo, RBD was selected as immunogen, rather than full-length S protein, inactivated SARS-CoV-2 whole virus or virus-like particles. The SARS-CoV-2 S gene was obtained by de novo synthesis with codon optimization. RBD expression plasmid was constructed as described in **Fig 1A**. To get large amounts of RBD proteins, the plasmid was transfected into CHO cells followed by an 8-day high-density fermentation. RBD-Fc proteins secreted into medium were purified by affinity chromatography against Protein A. Considering that Fc tag may induce unexpected antibodies in vivo, we completely removed Fc by thrombin digestion and conducting repeated purification against Protein A to remove residual Fc. Based on this method, gram-level RBD proteins were obtained.

**Figure 1.**
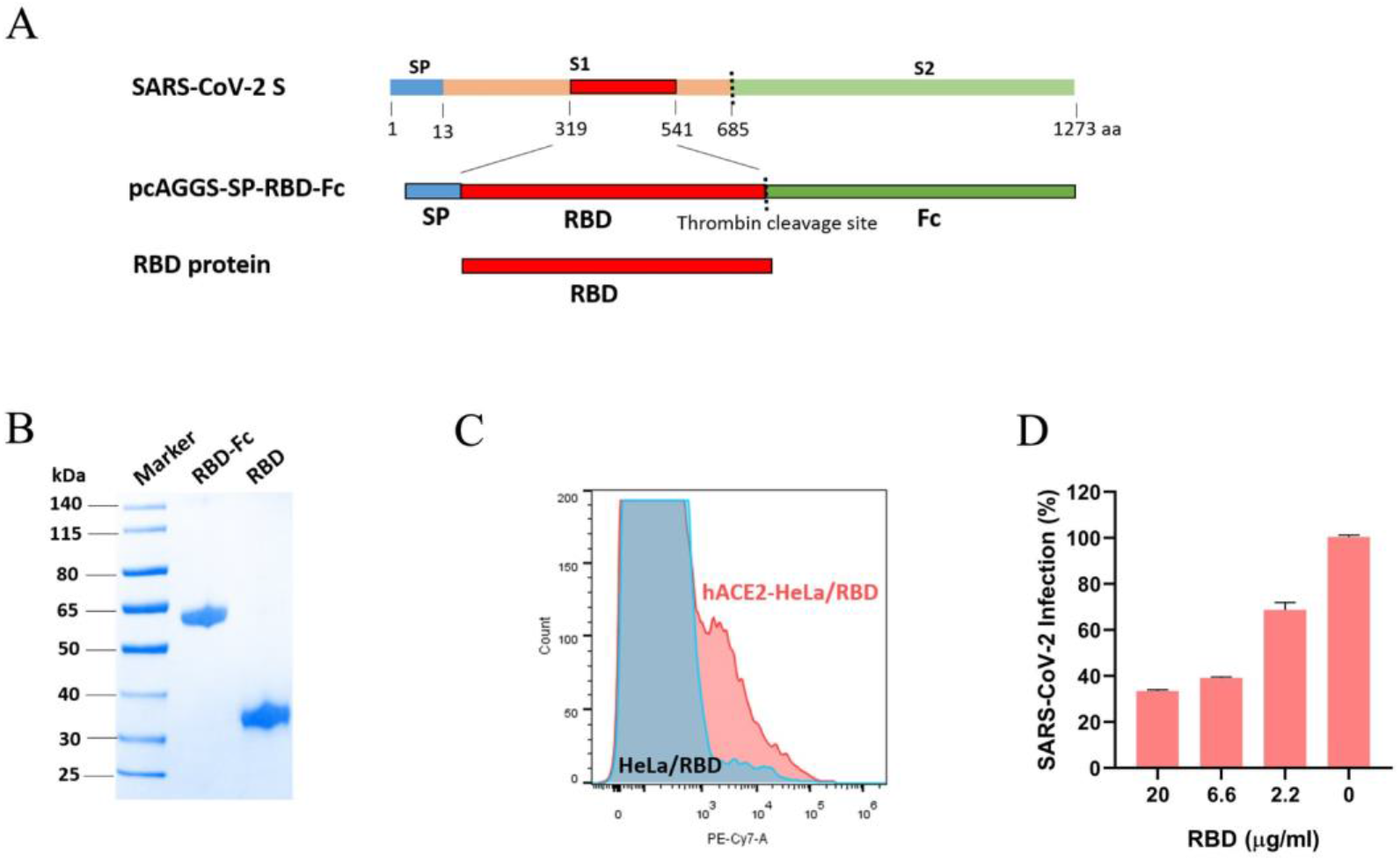
Preparation and characterization of RBD-Fc and RBD. **A** SARS-CoV-2 S protein contains a signal peptide, a receptor binding subunit S1, and a fusion subunit S2. The receptor binding domain (RBD) was predicted as 319-541 aa of S protein. Construction of RBD expression plasmid: RBD was constructed after an efficient signal peptide and followed by the thrombin site and linked with Fc as purification tag. RBD was obtained by CHO expression and purified from cell culture supernatant through affinity chromatography against Protein A, followed by thrombin digestion. **B** RBD-Fc and RBD were detected by reducing SDS-PAGE with Coomassie bright blue staining. Their sizes and purities are shown. **C** The binding of RBD to human ACE2 were detected by flow cytometry. HeLa cells were transfected with human ACE2 plasmid for 24 hours. Biotin-labelled RBD were incubated with the cells and followed by staining with fluorescent antibodies. The binding of RBD to ACE2-HeLa cells were observed as additional peaks compared to the HeLa cells. **D** Vero-E6 cells were pretreated with RBD at concentrations as shown in figure, then infected by SARS-CoV-2 at MOI=0.05. The blockade by proteins were calculated from the mock.

To examine the validity of RBD produced by our study, we characterized it by size, purity, and binding capacity to SARS-CoV-2 receptor, human ACE2. In **Fig 1B**, single strip was observed in the lane of RBD-Fc or RBD under the detection of reducing SDS-PAGE, implying high purities of these recombinant proteins. The binding activity of RBD to human ACE2 which was overexpressed on the surface of HeLa cells, was determined by flow cytometry. In contrast, RBD bound to Hela cells transiently transfected with human ACE2 plasmid, while rarely bound to HeLa cells with no ACE2 overexpression (**Fig 1C**). Furthermore, through cellular receptor blocking experiment, RBD inhibited the entry of SARS-CoV-2 in a dose dependent manner (**Fig 1D**). These results demonstrate a structural validity of RBD prepared in our study, guaranteeing its availability for further research.

### 2.2 Antigenicity and efficacy of SARS-CoV-2 RBD in mice

RBD was first tested in mice to examine their effectiveness in triggering antibody response in vivo. According a traditional immunization scheme described in Methods section, mice were immunized with 25 μg each mouse via subcutaneous injections with Freund’s adjuvant. As shown in **Fig 2A**, mice were immunized three times in total with two-week intervals, and sera samples were adopted ten days after each immunization to monitor the antibody response.

**Figure 2.**
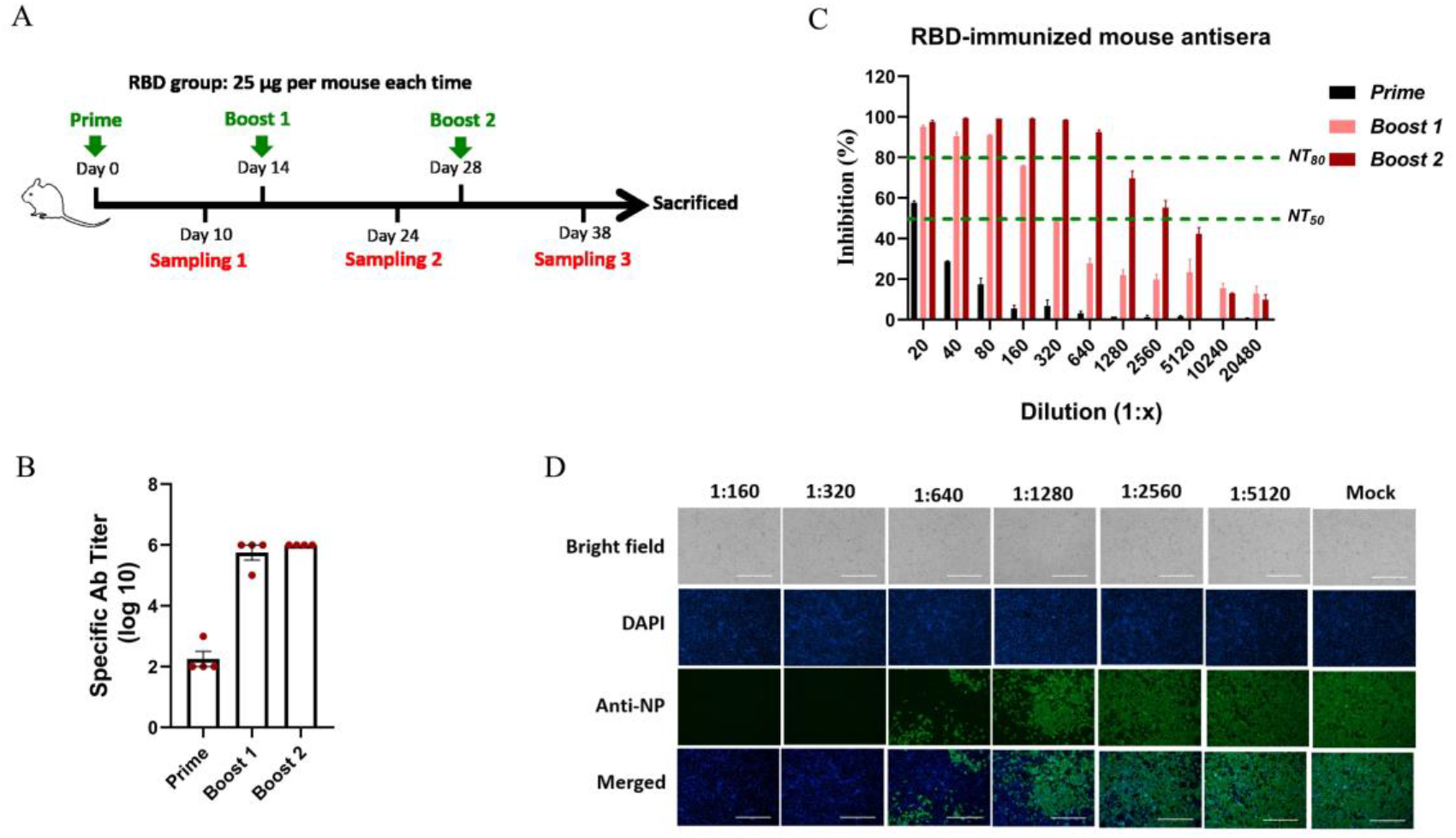
Antigenicity and efficacy of RBD in mice. **A** The scheme of mice immunization. Mice were immunized with 25 μg RBD per mice each time. The timeline of immunization and sampling were shown, and detailed immunization methods were described in the Methods section. **B** The titers of specific antibody in sera from RBD-immunized mice were detected by antigen-captured ELISAs. **C** NT_50_s and NT_80_s of sera from RBD-immunized mice were detected by a set of neutralization tests against live SARS-CoV-2 on Vero-E6 cells. Sera dilutions were from 1:20 to 1:20480 as shown in figures. NT_50_ and NT_80_ were marked with green lines. **D** The inhibition of sera from RBD-immunized mice after the third immunization was detected by IFA. Sera diluted as 1:160 to 1:5120 were used in neutralization tests, the infection of SARS-CoV-2 on Vero-E6 cells was detected by antibodies targeting NP. Cell nucleuses were stained with DAPI. Bright field and merged ones were also included.

In **Fig 2B**, the titers of specific antibodies for RBD were detected by antigen-captured ELISAs, and the titers were elevated with immunization times and reached 10^6^ after the third immunization, reflecting that RBD effectively elicit antibodies in mice. By neutralization test on SARS-CoV-2, sera from RBD–immunized mice after the second immunization inhibited 50% SARS-CoV-2 at a dilution of 1:320, and sera after the third immunization inhibited 50% SARS-CoV-2 at a dilution of over 1:2560, showing an immunization times-dependence (**Fig 2C**). And, NT_80_s of sera from RBD-immunized mice after the second immunization and third immunization achieved over 80 and over 640, respectively. The inhibition on SARS-CoV-2 by sera from RBD-immunized mice after the third immunization was also confirmed by indirect immunofluorescence analysis. The infection of SARS-CoV-2 on Vero-E6 cells was sharply reduced with the decrease of sera dilutions as shown in **Fig 2D**, showing a similar tendency with that in **Fig 2C**. These results collectively demonstrated that RBD could be used as immunogen in triggering nAbs in vivo.

### 2.3 Horse immunization and antisera production

Based on the above, RBD was taken as immunogen to produce horse antisera. Horses were immunized with RBD with complete Freund’s adjuvant at the first time and with incomplete Freund’s adjuvant at subsequent times, via intramuscular injections. The amount of RBD was doubled for the first three times, from 3 mg to 12 mg each horse, and fixed as 12 mg per horse when boosting before each plasma collection (**Fig 3A)**. Sera were adopted routinely after each immunization to monitor the antibody response.

**Figure 3.**
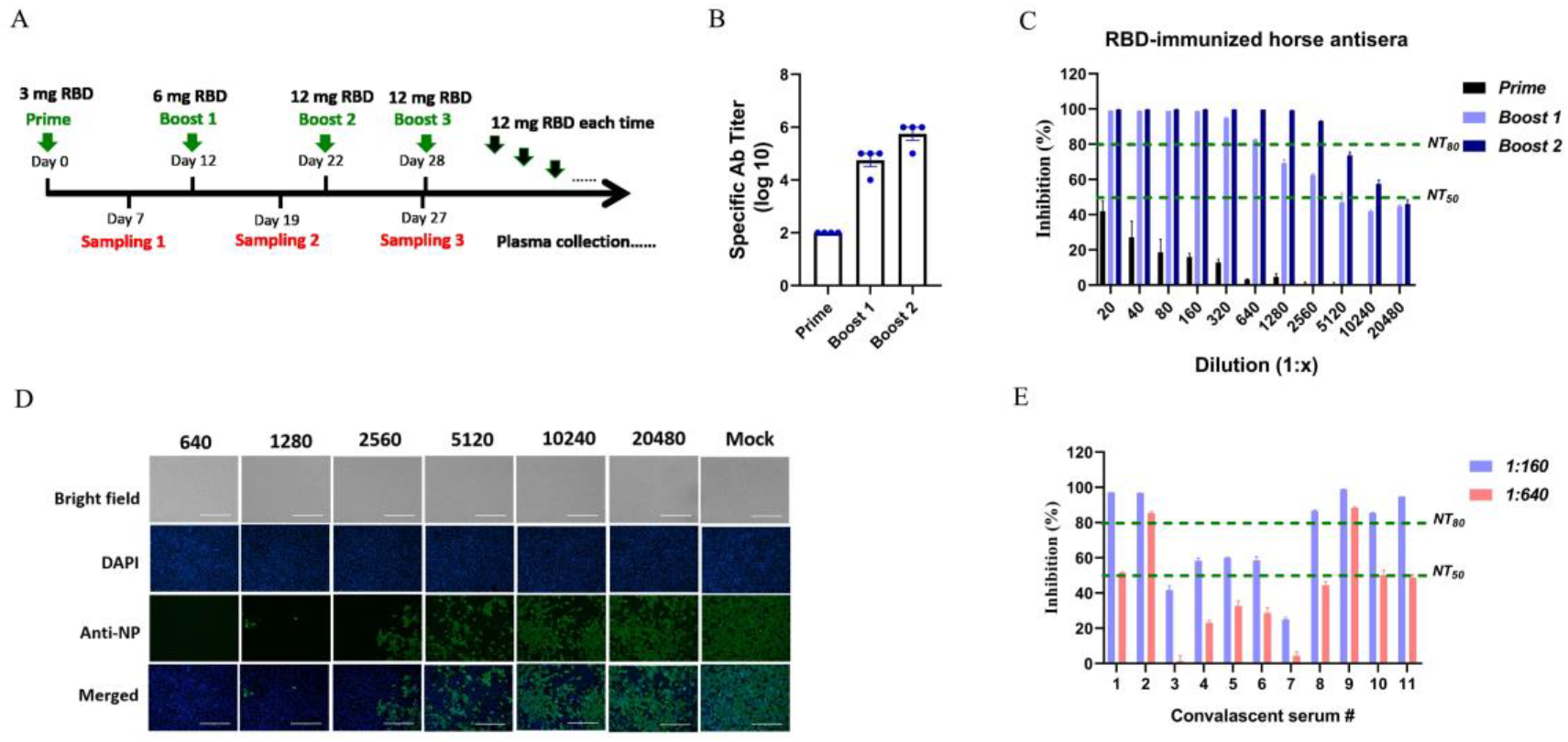
Horse immunization strategy and characterization of antisera against SARS-CoV-2. **A** The scheme of horse immunization. RBD was made as immunogen. Horses were vaccinated with 3 mg RBD on the first time with Freund’ complete adjuvant via intramuscular injections, then boosted with 6 mg RBD and 12 mg RBD on the second and third immunization with Freund’ incomplete adjuvant. Venous blood was adopted 7 days after each immunization for monitoring antibody response. Before each plasma collection, 12 mg RBD was immunized as usual. **B** The titers of specific antibody in horse sera after each immunization were examined by RBD-captured ELISAs. **C** NT_50_s and NT_80_s of sera from RBD-immunized horses were examined by neutralization tests against live SARS-CoV-2 on Vero-E6 cells. Sera dilutions were from 1:20 to 1:20480 as shown in figure. NT_50_ and NT_80_ were marked with green lines. **D** The inhibition of sera from RBD-immunized horses after the third immunization were detected by IFA. Sera diluted as 1:640 to 1:20480 was used in neutralization test, the infection of SARS-CoV-2 on Vero-E6 cells was detected by antibodies targeting NP. Cell nucleuses were stained with DAPI. Bright field and merged ones were also included. **E** The inhibition on SARS-CoV-2 by convalescent plasma was detected by a neutralization test on Vero-E6 cells as usual. Plasma was randomly adopted from 11 convalescent patients in Wuhan. The plasma were diluted as 1:160 or 1:640 as shown in figure.

As a consequence, the titers of specific antibodies were increased with immunization times and reached ~10^6^ after the third immunization (**Fig 3B**). NT_50_s of sera after the second and third immunization were 5120 and over 10240, and NT_80_s were 640 and over 2560, putting up high neutralization on SARS-CoV-2 (**Fig 3C**). Corresponding to that, the infection of SARS-CoV-2 on Vero-E6 cells was less than 20% under the treatment of sera after the third immunization with a dilution of 1:2560, the infection was less than 50% under the treatment with sera at a dilution of 10240 (**Fig 3D**). With the immunization strategy described in this study, high-titer neutralizing sera were obtained and made available for antibody preparation.

In comparison, 11 plasma samples, randomly adopted from patients in Wuhan, China, who recovered from COVID-19 in February 2020, were tested the same way as horse antisera. The results were, 5 out of 11 plasma samples (45%) inhibited over 50% SARS-CoV-2 with a dilution at 1:640, 2 out of 11 plasma samples (9%) inhibited over 80% SARS-CoV-2 with a dilution at 1:640, and 6 out of 11 plasma samples (55%) inhibited 80% SARS-CoV-2 needed a dilution of 1:160 (**Fig 3E**). While horse antisera after the third immunization inhibited 80% SARS-CoV-2 at a dilution of 1:2560 (**Fig 3C**). These demonstrated that passively immunized horses with RBD is more efficient in producing nAbs than purifying antibodies from convalescent plasma after natural infection by SARS-CoV-2, implying a necessity of producing horse antisera-derived antibodies.

### 2.4 Characterization of F(ab’)_2_ in vitro

Through pepsin digestion and purification described in the Methods section, F(ab’)_2_ was obtained from horse antisera. By a set of neutralization tests, F(ab’)_2_ was found inhibiting SARS-CoV-2 with EC_50_ as 8.78 μg/ml and EC_80_ as 24.92 μg/ml (**Fig 4A**). As shown in **Fig 4B**, over 90% SARS-CoV-2 were inhibited under the treatment of F(ab’)_2_ at 31.15 μg/ml, and over 50% SARS-CoV-2 were inhibited at 7.81 μg/ml, the inhibition on SARS-CoV-2 was observed with apparent dose-dependent manner. Furthermore, the kinetics of binding to and dissociating from recombinant RBD were determined by biomolecular interaction analysis, and the *K*_*D*_ of total F(ab’)_2_ to RBD was 75.6 nM (**Fig 4C**).

**Figure 4.**
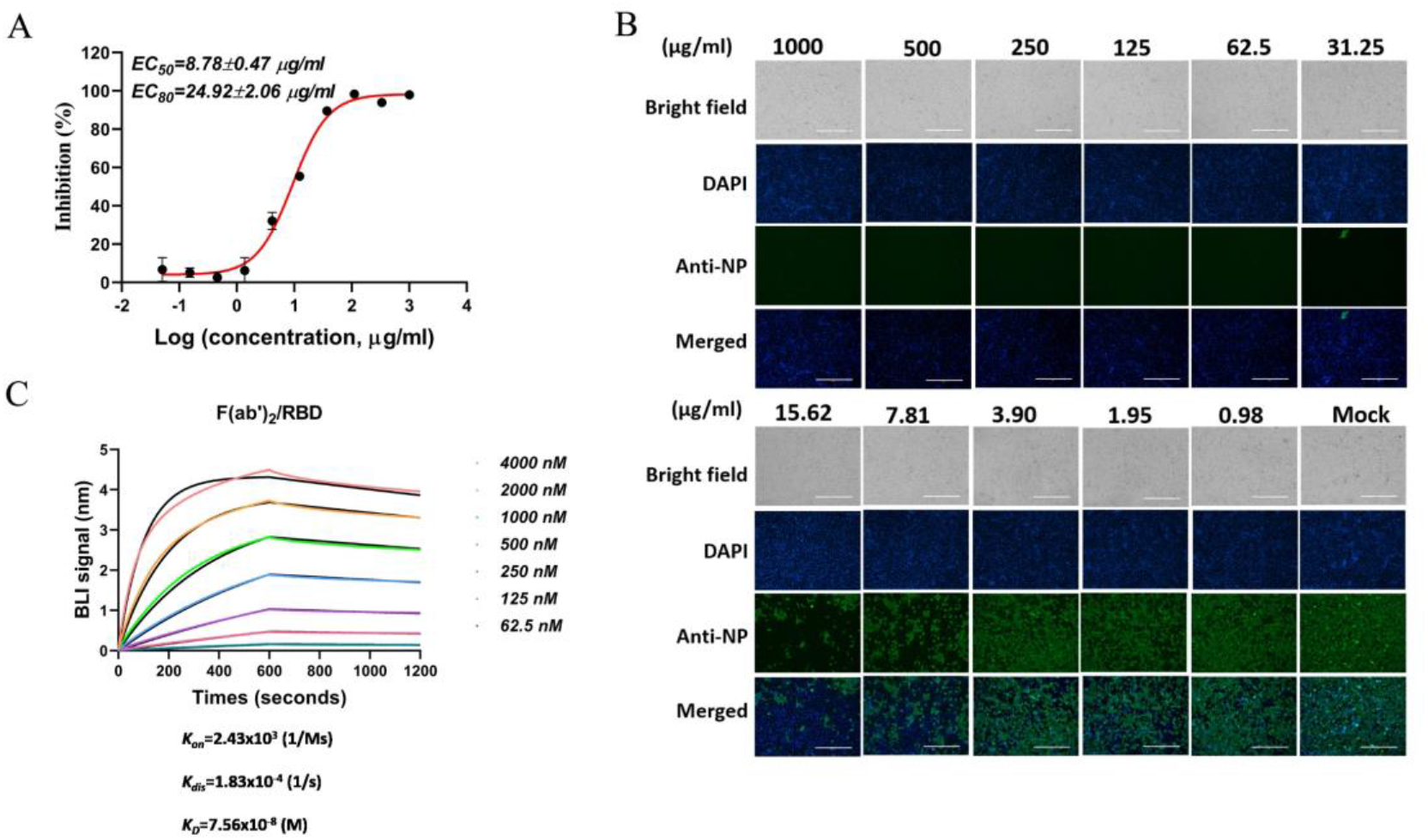
Neutralization of F(ab’)_2_ on SARS-CoV-2 and the binding of F(ab’)_2_ to RBD. **A** Inhibition of F(ab’)_2_ on SARS-CoV-2 was examined by a neutralization test as usual, EC_50_ and EC_80_ were calculated as 8.78 and 24.92 μg/ml, respectively. **B** Infection of SARS-CoV-2 on Vero-E6 cells under the treatment of F(ab’)_2_ was detected by IFA against NP. Working concentrations of F(ab’)_2_ were from 1000 to 0.98 μg/ml. **C** Affinity of F(ab’)_2_ to SARS-CoV-2 RBD was detected by BLI, and the kinetics (*K*_on_ and *K*_dis_) were processed by an Octet data analysis system. The binding curves were obtained by passing F(ab’)_2_ at concentrations from 4000 to 62.5 nM over biotinylated RBD immobilized on a streptavidin biosensor surface. The kinetic values *K*_D_, M) were calculated by deducting from baseline and fitting the association and dissociation responses to a 1:1 Langmuir binding model.

To further improve the neutralizing activity of F(ab’)_2_, high binders to RBD were purified from total F(ab’)_2_ by affinity chromatography against RBD. The neutralizing activity of RBD-specific F(ab’)_2_ was tested the same way as total F(ab’)_2_. As expected, RBD-specific F(ab’)_2_ neutralized SARS-CoV-2 with EC_50_ as 0.07μg/ml and EC_80_ as 0.18 μg/ml, respectively (**Fig 5A**), showing a strong inhibitory effect on SARS-CoV-2. Correspondingly, the high affinity of RBD-specific F(ab’)_2_ to RBD is reflected by a *K*_*D*_ as 0.76 nM (**Fig 5B**). The correlation between neutralizing activity on SARS-CoV-2 and affinity to RBD suggested that F(ab’)_2_ produced by our study targets RBD to work. The potent neutralization potentiates F(ab’)_2_ as an alternative for SARS-CoV-2.

**Figure 5.**
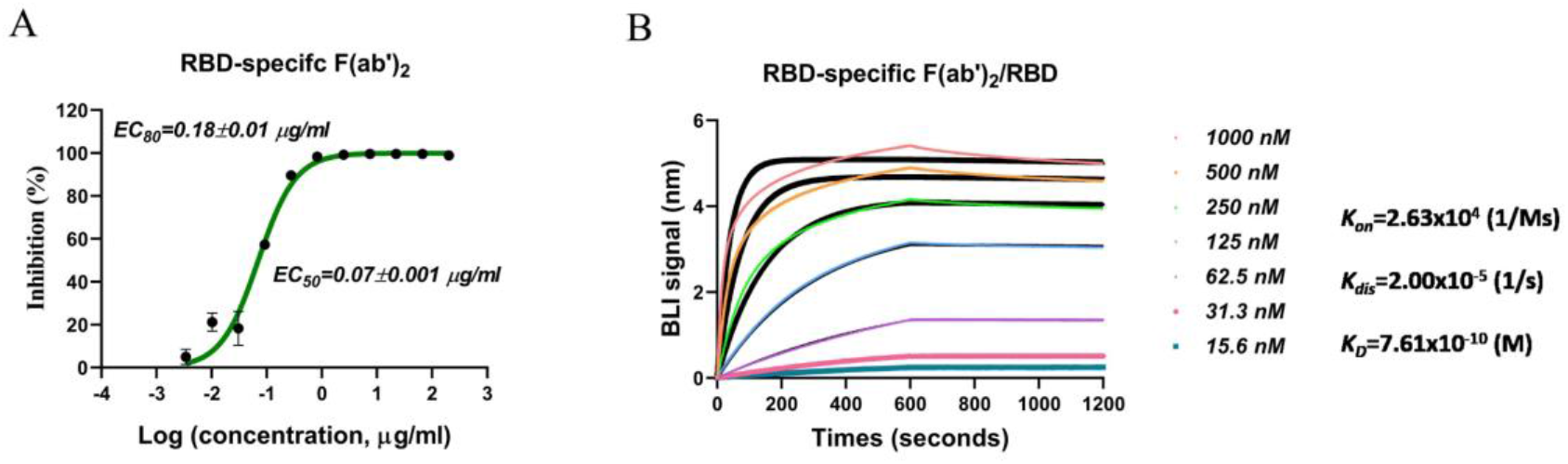
Neutralization of RBD-specific F(ab’)_2_ on SARS-CoV-2 and the binding of RBD-specific F(ab’)_2_ to RBD. **A** Inhibition of RBD-specific F(ab’)_2_ on SARS-CoV-2 was examined by neutralization test as usual, EC_50_ and EC_80_ were calculated as 0.07 and 0.18 μg/ml, respectively. **B** Affinity of RBD-specific F(ab’)_2_ to recombinant RBD was detected by BLI as described in Figure 4c legend. The binding curves were obtained by passing F(ab’)_2_ at concentration from 1000 to 15.6 nM over biotinylated RBD immobilized on a streptavidin biosensor surface.

## 3 Materials and Methods

### 3.1 Cells lines and viruses

Vero E6 and HeLa cells were maintained in Dulbecco’s Modified Eagle’s Medium (DMEM, Gibco, Grand Island, NY) supplemented with 10% fetal bovine serum (FBS, Gibco) at 37°C with 5% CO_2_. ExpiCHO-S cells were cultured with ExpiCHO Expression Medium (Gibco) in an incubator at 37°C and 8% CO_2_ while shaking at 125 rpm/min.

SARS-CoV-2 live viruses were from National Virus Resource, Wuhan institute of Virology, Chinese Academy of Sciences, and handled in BSL-3 lab. SARS-CoV-2 viruses were passaged on Vero-E6 cells.

### 3.2 Protein expression and purification

The gene of SARS-CoV-2 S RBD (319-541 aa) was synthesized in GenScript Co., Ltd., and cloned to eukaryotic expression plasmid pCAGGS to obtain pCAGGS-Signal peptide-RBD-(Thrombin site)-Fc. The plasmid was purified from *E. coli* (DH5α) with an endotoxin-free plasmid extraction kit (Invitrogen, Carlsbad, CA) and transfected into ExpiCHO-S cells using ExpiFectamine Transfection Kit (Gibco). Cell supernatants containing RBD-Fc were collected 8 days later and filtrated with 0.22 μm film before being subjected to affinity chromatography through Protein A agarose. The eluted fraction containing RBD-Fc was taken for further analysis. RBD without Fc tag was obtained from RBD-Fc by removing Fc through thrombin digestion. RBD proteins were collected from flow-through faction, affinity chromatography against Protein A were repeatedly conduced to completely remove residual Fc.

### 3.3 Reducing SDS-PAGE

RBD-Fc and RBD were analyzed by SDS-PAGE with Coomassie brilliant blue staining. To verify the sizes and purities of RBD-Fc and RBD prepared in our study, reducing SDS-PAGE were employed. Reduced samples were prepared by mixing 2 μg proteins with loading buffer, adding 2-β-mercaptoethanol, and then boiling in water for 10 minutes. The samples were concentrated with 4% SDS-PAGE and separated with 10% SDS-PAGE. Finally, SDS-PAGE gels were stained with Coomassie Brilliant Blue. An image of the gel after decolourization was captured with a ChemiDoc MP Imaging system (Bio-Rad).

### 3.4 Flow cytometry

The binding of RBD to human ACE2 was detected by flow cytometry as described elsewhere^23^. HeLa cells were seeded in 6-well plate overnight. PcDNA3.1-human ACE2 was transfected into HeLa cells with Lipo2000 (Invitrogen), PcDNA3.1 was made as control. 24 hours later, cells were scraped off and washed with PBS. RBD was labelled with biotin and dialyzed with PBS to remove residual biotin before use. Biotin labelled-RBD was added to ACE2-overexpressed cells or mock cells up to a final concentration of 10 μg/ml. After incubation at 4°C for 30 minutes, cells were washed with PBS. Then PE Cy7-conjugated streptavidin (BD bioscience, CA, USA) were added to cells. After incubation at 4°C for 15 minutes, cells were washed and suspended with PBS before detecting with cytometry (BD).

### 3.5 Receptor blocking assay

Inhibition of SARS-CoV-2 entry by RBD protein was carried out, as previously described, with some modifications^24^. Vero-E6 cells were seeded in 48-well plates with 5 × 104 cells/well overnight. After removing culture medium, RBD diluted in 2% FBS-DMEM as 20, 6.6 and 2.2 μg/ml were incubated with cells at 37°C for 1 hour, PBS was made as control. After that, proteins were removed, cells were washed twice with PBS. SARS-CoV-2 with MOI=0.05 in 100 μl 2% FBS-DMEM were added to each well. After incubation at 37°C for 1 hour, supernatant was completely removed, cells were washed twice with PBS before adding fresh 2% FBS-DMEM. 24 hours later, cell supernatant was collected for viral copy detection. Infection (%) of SARS-CoV-2 was calculated from control.

### 3.6 Mouse immunization and sampling

Female BALB/c mice aged 6-8 weeks were housed in specific pathogen-free animal care facilities. According to a homogeneous prime-boost-boost protocol, immunization was performed three times in total with two-week intervals. In detail, 25 μg RBD in a volume of 100 μl PBS were mixed with 100 μl Freund’s complete adjuvant (Sigma-Aldrich, St. Louis, MO) for priming or mixed with 100 μl Freund’s incomplete adjuvant (Sigma-Aldrich) for boosting. A total of 200 μl mixture were used for subcutaneous injections in each mouse. Blood samples were adopted from ophthalmic vein 10 days after each immunization.

### 3.7 Horse immunization and sampling

Four healthy horses aged 6-10 years old after quarantine inspection were housed in standard breeding conditions. Before immunization, 3 mg RBD protein in 3 mL PBS were mixed with equal-volume Freund’s complete adjuvant (Sigma-Aldrich) for priming, 6 mg RBD protein in 3 mL PBS were mixed with equal-volume Freund’s incomplete adjuvant (Sigma-Aldrich) for the first boosting, and 12 mg RBD protein in 3 mL PBS was mixed with equal-volume Freund’s incomplete adjuvant (Sigma-Aldrich) for the second boosting. Horses homogeneously received intramuscular injections at day 0, 12, and 22. Serum samples were collected from the jugular vein on days 7, 19, and 27 for monitoring the variation of antibody response. Large amount of plasma collection were collected after the third boosting and each collection was conducted 7 days after boosting with 12 mg RBD each horse.

### 3.8 Enzyme-linked immunosorbent assay (ELISA)

Recombinant RBD protein diluted at 2 μg/ml was coated on 96-well plate with 100 μl/well overnight at 4°C. The liquid was aspirated, and plate was washed three times with PBS-0.1% Tween 20 then blocked with 2% nonfat-milk at 37°C for 1 hour. Gradient diluted mouse or horse sera were added to each well and incubated at room temperature for 1 hour. PBS and irrelevant sera were used as controls. Then, the liquid was aspirated, and the plate was washed five times with PBS-0.05% Tween 20 and incubated with secondary antibodies conjugated with (HRP) at room temperature for 1 hour. After washing five times as usual, 3, 3’, 5, 5’-tetramethylbenzidine was added, and the chromogenic reaction was terminated by adding H_2_SO_4_ about 10 minutes later. Finally, the absorbance at 450 nm was measured with a microplate reader (TECAN, Swiss), and values greater than twice those of the controls were considered positive.

### 3.9 Virus neutralization test

Vero-E6 cells were seeded in 48-well plates with 5 × 10^4^ cells/well overnight. Mouse or horse sera, human convalescent plasma or F(ab’)_2_, were firstly diluted in 100 μl 2% FBS-DMEM, and incubated with 5 μl SARS-CoV-2 (MOI=0.05) at 37 °C for 1 hour. Then cell supernatants were aspirated, 100 μl antisera- or antibody-virus mixture were added. After incubation at 37 °C for 1 hour, the supernatant were completely removed, cells were washed with PBS and supplemented with fresh 10% FBS-DMEM. After 24 hours, cell supernatants were collected and subjected to viral RNA isolation and cells were kept for indirect immunofluorescence analysis. Viral genome copies were detected by qRT-PCR with primers targeting S gene.

### 3.10 Indirect immunofluorescence assay (IFA)

Cell plates were collected after a virus neutralization test. Cells were washed with PBS then fixed with 4% paraformaldehyde and permeabilized with Triton x-100. After that, cells were blocked with 2% nonfat-milk at room temperature for 1 hour, then were washed with PBS and incubated with rabbit anti-NP antibodies at room temperature for 2 hours. Cells were washed again before incubation with goat anti-rabbit Alexa Fluor 488-conjugated antibodies at room temperature for 1 hour. Finally, cells were washed and stained with 4, 6-diamino-2-phenylindole (DAPI) for 10 minutes at room temperature. Images were captured by a fluorescence microscope (Olympus, Japan).

### 3.11 Preparation of F(ab’)_2_

When neutralizing titer of horse antisera met requirement as NT_50_ over 10000, the plasma were collected with the use of a plasma collection machine 7 days after each boosting. Briefly, plasma was firstly diluted with bi-distilled water at a ratio of 1:4 and the pH was adjusted to 3.0 with HCl. Then pepsin (Sigma) was added and the temperature was adjusted to 30°C. The incubation was preserved for 1.5 hour with stirring. Pepsin was inactivated by temperature evaluation, then ammonium sulfate with gradient concentration was added in proper order, each followed by filtration. Finally, the supernatant was subjected to a Protein A column to remove residual IgG, and F(ab’)_2_ were collected from flow-through fraction.

### 3.12 Biomolecular interaction analysis (BIA)

The affinities of F(ab’)_2_ to RBD were monitored by biolayer interferometry (BLI) using an Octet-Red 96 device (Pall ForteBio LLC., CA) according to previously described protocols ^25^. Briefly, RBD was biotinylated at room temperature for 0.5 hours by incubating with biotin at a molar ratio of 1:3. Residual biotin was removed by dialysis with PBS. Biotinylated RBD at 10 μg/ml was loaded onto streptavidin biosensors (ForteBio) until saturation, and F(ab’)_2_ or RBD-specific F(ab’)_2_ diluted to 4000, 2000, 1000, 500, 250, 125, 62.5 nM or 1000, 500, 250, 125, 62.5, 31.3, 15.6 nM were then loaded. The kinetics of association (*K*_on_) and dissociation (*K*_dis_) were measured, the data was processed by an Octet data analysis system.

### 3.13 Purification of RBD-Specific F(ab’)_2_

For purification of RBD-specific F(ab’)_2_, RBD protein expressed by CHO cells and prepared as described above, was coupled on pre-activated resin (PabPurSulfolink Beads, SMART Life Sciences, Changzhou) through amino reaction^17^. And the RBD-coupled resin was then used to purify RBD-specific F(ab’)_2_ from total F(ab’)_2_. In brief, total F(ab’)_2_ were diluted in 20 mM phosphate buffer (pH 8.0) and repeatedly flowed through RBD-coupled resin to make binding to RBD. Then resin was adequately washed with 20 mM phosphate buffer (pH 8.0) before adding Glycine (1 M) to elute high binders. The eluted component, RBD-specific F(ab’)_2_, was dialyzed with PBS to remove glycine and maintained in PBS before use.

### 3.14 Ethics statements

All animal experiments were performed strictly according to the Regulations for the Administration of Affairs Concerning Experimental Animals in China, and the protocols were approved by the Laboratory Animal Care and Use Committee of Wuhan Institute of Virology, Chinese Academy of Sciences (Wuhan, China).

### 3.15 Data analysis

Data was analyzed using GraphPad Prism 8.0 software (San Diego, CA, USA), are presented as mean±SD based on at least three independent experiments.

## 4 Discussion

COVID-19 is an emerging infectious disease caused by a new member of the coronavirus family. SARS-CoV-2 became a serious threat to global public health within a short time period during the outbreak. Although biotechnology and pharmaceutical technology has grown rapidly in the 21st century, humans remain passive in responding to public health emergencies. Due to the close contact with wild animals such as pangolin and bats^23–25^, infectious diseases caused by coronaviruses such as SARS- and MERS-CoV along with SARS-CoV-2 uninterruptedly pose threats to mankind. This is a reminder of developing broad-spectrum drugs and vaccines^26^, when the pace of specific drug development is not fast enough to cope with. Prior experience in infectious disease control suggests that antiviral serum obtained from hyper immune equine plasma has long been used for the treatment of life threatening viral diseases^27^. We showed that horse immunoglobulin fragment F(ab’)_2_ has a potential to provide protection for COVID-19.

Clinical evidence showed that the latent period of COVID-19 is short (about 5 days to 2 weeks) and that most patients appear to recover within a short time with no persistent or latent infection in the organism, it is reasonable to conclude that a neutralizing antibody may play an important role in preventing SARS-CoV-2 infection^28^. Shen et *al*. recently reported that administration of convalescent plasma containing neutralizing antibody could improve COVID-19 patient clinical status^13^. Convalescent plasma from patients has played important roles in the treatment of infectious diseases^29^. The convalescent plasma was used for severe acute respiratory syndrome (SARS)^30, 31^, H5N1 influenza infection^32, 33^, Ebola^34^, and other viral infections, while it has the risk of blood-borne disease and usually suffers from insufficient blood sources.

Seventeen years ago, we prepared inactivated SARS-CoV whole viruses and the test in mice proved that the inactivated vaccine induced specific antibodies to SARS-CoV, and a neutralization test in vitro also proved that the induced antibodies could neutralize SARS-CoV^35^. A major problem meriting further study in the induction of neutralizing antibodies by an inactivated SARS-CoV whole virus is that some neutralizing antibodies may enhance the infection of the coronavirus^36^. Several studies showed antibody-dependent enhancement (ADE) of SARS coronavirus infection^37, 38^. Additionally, reports of antibodies induced by the enhanced infection are found in studies of HIV, SIV and Dengue virus^39–41^. Thus, the SARS-CoV-2 spike protein receptor binding domain (RBD) was selected as immunogen in this study. It could induce highly effective neutralizing antibodies with higher safety.

In addition, RBD-specific horse F(ab’)_2_, as a relatively specific antibodies, has its unique advantages in contrast to monoclonal antibodies and human immunoglobulin after natural infection. It specifically targeted RBD, prevented the binding of virus to its receptor, ACE2, bringing no risk associated with Fc, further avoiding ADE. Meanwhile, it binds multiple epitopes of SARS-CoV-2 RBD, hinting a broad-spectrum neutralization on human or bat SARS-related coronaviruses and bat SARS-like coronaviruses. Furthermore, we demonstrated that neutralizing titer of horse antisera by immunization with RBD is higher than that of convalescent human plasma after natural infection by SARS-CoV-2 in this study. Moreover, the horses could be immunized more times. Higher titer of therapeutic antibodies for SARS-CoV-2 obtained from hyper immune equine plasma was expected.

In this study, large amounts of horse antisera were prepared by immunization with SARS-CoV-2 RBD. Before making RBD as immunogen, we verified its conformation by receptor binding experiments and tested its antigenicity in mice. According a strategy of immunogen dose-increasing strategy, high-titer horse antisera were obtained, and F(ab’)_2_ were manufactured in a GMP workshop to be processed for clinical study. As a matter of fact, monoclonal antibodies against SARS-CoV, like CR3022 potently binds SARS-CoV-2 while its neutralization on SARS-CoV-2 has not been verified^42^. Other neutralizing antibodies against SARS-CoV were proved weakly binding SARS-CoV-2 RBD^43^. F(ab’)_2_ prepared by our study can potentially neutralize SARS-CoV to some extent, since it contains antibodies binding multiple epitopes on RBD, in addition to that amino acid sequence identity between SARS-CoV-2 and SARS-CoV RBDs is 73% and these two employed the same receptor ACE2 to enter host cells.

Besides, F(ab’)_2_ is a kind of immunoglobulin fragment, which is prepared by removing Fc from IgG and retaining the active fragment, significantly reducing side effects in the body. Although horse F(ab’)_2_ is a heterologous protein to human immune system, horse serum proteins and Fc-related proteins are removed as much as possible using modern industrial techniques. Also, single-dose administration will avoid the adverse effects and efficacy falling induced by repeated administration. Additionally, the operation of Fc removal eliminates the major concern about ADE in coronaviruses. These enable antibody drugs such as horse F(ab’)_2_ to be candidates for COVID-19 therapy.

## 5 Conclusions

In summary, we herein successfully obtained therapeutic antibodies from hyper immune equine plasma. Horse immunoglobulin fragment F(ab’)_2_ against RBD was highlighted as a potential therapeutic for COVID-19.

## 6 Acknowledgements

This study was financially supported by grants from the National ministry of science and technology emergency project (2020YFC0841400) and the Hubei provincial department of science and technology emergency project (2020FCA003). This study was also supported by the Wuhan National Biosafety Laboratory of the Chinese Academy of Sciences.

We would like to thank Ding Gao and Juan Min at the Centre for Instrumental Analysis and Metrology, Wuhan Institute of Virology, Chinese Academy of Sciences for providing technical assistance. We also thank P3-lab staff in Wuhan Institute of Virology, Chinese Academy of Sciences.

## 7 Conflict of Interest

The authors declare no competing financial interests.

## 8 Authors’ Contributions

X.-Y. P. drafted the manuscript, processed data, and did partial mouse and horse experiments. P.-F. Z., J. Z., L.-J. F. and J. S. prepared immunogen. Y. W., W.-J. S., X.-M. J. and Y. S. did neutralization tests and viral detection. T.-J. F. and X.-Y. S. vaccinated horses and prepared F(ab’)_2_. S.-J. Z. and R. G. constructed plasmid. Z. C. and G.-F. X. revised the manuscript and supervised the project process.

